# The biotoxin BMAA promotes mesenchymal transition in neuroblastoma cells

**DOI:** 10.1101/2024.11.07.622435

**Authors:** Ellie Moore, Abby Renner, Taylor Dowden, Devin Messer, Hayes Hansen, McKinnely Mull, Abigail Goebel, Grace Mawawa, Bryan Burton, Matthew Pawlus

## Abstract

Mesenchymal-like cancer cells are an indicator of malignant tumors as they exhibit tumorigenic properties including downregulation of differentiation markers, and increased colony-forming potential, motility, and chemoresistance. We have previously demonstrated that the cyanobacterial biotoxin beta-methylamino-L-alanine (BMAA) is capable of influencing neural cell differentiation state through mechanisms involving the Wnt signaling pathway, suggesting the possibility that BMAA may play a role in influencing other Wnt related differentiation processes including mesenchymal transition. In this study we present evidence characterizing the effects of BMAA on mesenchymal transition in a human neuroblastoma cell line and provide support for the hypothesis that the biotoxin can promote this process in these cells by altering differentiation state, inducing changes in gene expression, and changing cellular function in manners consistent with cellular mesenchymal transition. Results of this study indicate that BMAA exposure may promote carcinogenesis through its effects on cell differentiation state in certain contexts. These results suggest that exposure to the biotoxin BMAA may be an influencing factor in chemotherapy resistance and cancer relapse in neuroblastoma.

## Introduction

Neuroblastoma (NB) is the most frequently occurring extracranial tumor for children, arising from neural crest progenitor cells during development [1]. This disease is characterized by its variability in presentation making it difficult to treat [2]. While many low-risk patients require little to no treatment, almost half of children diagnosed are considered high-risk and require multimodal therapies. Among those that do reach remission post-therapy, 40-50% will suffer a malignant relapse due to the cancer’s aggressive nature and resistance to current options [3]. Evidence suggests that cancer cell de-differentiation to a mesenchymal-like state may promote the occurrence of relapse in patients [4]. Therefore, gaining a more complete understanding of the factors influencing cancer cell differentiation state may prove beneficial to understanding and treating neuroblastoma.

Epithelial-to-Mesenchymal Transition (EMT) in cancer cells is characterized by cell de-differentiation from the epithelial state to a mesenchymal cell-like phenotype [4]. Mesenchymal-like cancer cells possess a set of specific characteristics that promote cancer progression and relapse including increased cell motility, single cell survival, proliferation, and chemoresistance [5,6]. This transition is induced by various factors including hypoxia, cytokines, antitumor drug treatment, and growth factors produced in a tumor environment [6,7]. These factors are known to promote activation of developmental signaling pathways such as epidermal growth factor (EGF)/epidermal growth factor receptor (EGFR), hedgehog, Notch, tumor growth factor-β (TGF-β), and Wnt/β-catenin, which play a role in development and EMT [8]. Numerous studies suggest that inhibiting Wnt can inhibit EMT in both in human cells as well as animal studies [9, 10]. Wnt inhibitors are also clinically used as treatment for other cancers [10].

A cyanobacterial biotoxin, beta-methylamino-L-alanine (BMAA), has previously been demonstrated to upregulate Wnt signaling in a neuroblastoma cell line and potentially affect neural cell differentiation state [11]. BMAA is an environmental toxin produced by cyanobacteria and eukaryotic diatoms within algae blooms of freshwater and marine environments and has been associated with neurodegeneration in numerous studies [12]. Its role in other contexts like cancer, however, have not been extensively investigated. Human exposure occurs through potential food sources correlated with water ecosystems hosting BMAA-producing microorganisms. Prior studies have demonstrated the potential for ingestion of environmental contaminants such as N-nitroso compounds in water sources to contribute to the development of neural cancers arising from the central nervous system [13, 14]. Thus, we propose that exposure to BMAA may influence neuroblastoma development, specifically by promoting EMT through Wnt activation. As temperatures continue to rise, so will the presence of large algal blooms capable of producing this biotoxin and potential human exposure [15]. The worldwide presence of BMAA coupled with accelerated climate change both highlight the importance of elucidating the role of BMAA in human health contexts.

## Materials and Methods

### Cell culture

IMR-32 cells (ATCC, Manassas, VA) were cultured in EMEM (Gibco, Grand Island, NY) supplemented with 10% FBS (Gibco). This cell line was incubated under standard growth conditions of 37° C and 5% carbon dioxide. Cells grown in a monolayer were cultured on a flat-bottomed plate, while tumorspheres were cultured on an ultra-low attachment round-bottomed plate.

### Microscopy and imaging

Cells were observed either in culturing vessels or fixed on slides using an AmScope LB-702 fluorescent microscope (Irvine, CA) under the indicated magnification. Images were captured using the AmScopeAmLite 20200526 microscopy software and analyzed using ImageJ software where appropriate.

### Gene Expression

RNA was isolated from cells using the RNeasy Plus Micro Kit (Qiagen, Hilden, Germany), utilizing DNase to digest possible contaminated genomic DNA. RNA was reverse transcribed using the SuperScript IV Reverse Transcriptase (Invitrogen, Waltham, MA). Levels of mRNA were quantified using a TaqMan Human Molecular Mechanisms of Cancer Array (Applied Biosystems, Foster City, CA) that utilizes real-time qPCR. Results for the cancer genes were run through GO ontology database and a PANTHER overrepresentation test to observe pathway linkages. Relative cDNA abundance for Wnt gene expression was quantified using a TaqMan Human Wnt Pathway Array (Applied Biosystems, Foster City, CA) via probe hybridization and chemiluminescence detection.

### Clonogenic Assay

IMR32 cells were cultured as a 2D (monolayer) culture in a 12 well dish and treated with either 2uM or 4uM of BMAA. Media was removed 36 hours post-treatment and cells were stained with methionine blue for counting. Single cell survival was measured by manually counting colony abundance per well. Triplicates of each condition were counted and averaged to obtain an average colony count per 5K cell well.

### Scratch Assay

IMR-32 cells were cultured as a 2D (monolayer) culture for 48 hours in the presence of either DMSO, BMAA, or Chiron all with cell growth media. Cell media was removed from all wells and a scratch assay was implemented by running a 0.1-20uL micropipette tip in a straight line from the north to south end of each well. Width of the scratch was measured east to west from the outermost edge of the scratch using the straight-line tool in ImageJ. Cell motility was assessed by finding the average scratch width from triplicates for each condition prior to and after 48 hours with treatment. By comparing each condition at 48 hours and 0 hours respectively, average percent decrease in scratch width was found. A higher average percent decrease correlates with increased cell motility.

### Proliferation, Tumorspheres

IMR-32 cells were cultured under normal conditions as a monolayer for seven days to allow for tumorsphere growth. Tumorspheres were treated with various concentrations of BMAA (2 and 4uM) and incubated over a period of 36 hours. Triplicates for each condition (control, 2uM, and 4uM) were counted manually and averaged to obtain average sphere number. Sphere size was measured with the straight-line feature of the software tool ImageJ. Using this feature the longest cross width diameter of three spheres per well was measured and averaged to get average sphere size per well. This was performed for three wells per condition and triplicates were averaged to obtain average relative sphere size. Tumorspheres for the Myc inhibitor experiment followed the same monolayer culture protocol. Spheres formed on the monolayer were transferred individually into a 96 well round bottom dish using a 200-1000uL micropipette tip. Treatment with BMAA alone or co-treatment with BMAA and XAV939 (1uM) or KJ Pyr9 (100uM) was administered and tumorspheres were incubated over a period of seven days. At various time points from treatment (day 0, 3, 5, 7) photos were taken for triplicates of each condition and sphere diameter was assessed using the straight-line tool in the ImageJ analysis software. Triplicates for each condition were measured and averaged to obtain average sphere size per day measured.

### Cytotoxicity

IMR-32 cells were cultured as a monolayer or tumorsphere under normal conditions with increasing concentrations of BMAA (1, 2, 4uM) and Dox (0.002, 0.004, 0.008, 0.016, 0.032, 0.064, 0.128 uM), and incubated over a period of 36 hours. To assess cell death the presence of extracellular lactate dehydrogenase (LDH) was assessed in cell media using the CyQUANT LDH Cytotoxicity Assay Kit (Invitrogen, Waltham, MA) according to the manufacturer’s instructions. Cell lysis was quantified using a Molecular Devices SpectraMax Microplate Spectrophotometer (San Jose, CA).

### Matrigel Transwell Assay

IMR-32 cells were cultured on 12 inserts of a 24 well Corning BioCoat Matrigel Invasion Chamber pre-coated with an 8.0 µm PET Membrane. IMR-32 cells were treated with 4uM of BMAA, as well as 1uM XAV939, 100uM KJ Pyr9, and 1uM CHIR99021 and incubated over a period of 48 hours. Cells were serum starved by removing growth media and performing a wash with PBS. To quantify cell migration, Matrigel inserts were removed and the lower side of the transwell membrane was stained with crystal blue and quantified by manually counting the colonies.

### Statistics

Experimental data from a minimum of three replicate experiments for each assay were analyzed by 2-tailed student’s T-test unless otherwise noted and significant p-values reported in each figure; *p<0.05, **p<0.001 vs. untreated cell control; #p<0.05, ##p<0.001 vs. paired experimental sample.

## Results and Discussion

### Neuroblastoma

The relationship between neuroblastoma and the epithelial to mesenchymal transition pathway has been thoroughly investigated [16,17]. However, little is known about the role that BMAA and the Wnt pathway plays in this relationship. Obtaining a more complete understanding of the factors contributing to EMT may influence our knowledge and treatment of neuroblastoma.

### BMAA plays a role in cancer progression

The Wnt/ β-catenin signaling pathway is widely known as a primary regulator of cell development and differentiation state. Irregularities in this signaling pathway have been tied to cancer stem cell renewal, proliferation, and differentiation [18]. Prior studies show evidence that the toxin BMAA dysregulates Wnt signaling, but little is known about how exposure to the toxin may influence cancer development [11]. We selected IMR-32 human neuroblastoma cells as they exhibit a moderately aggressive neuroblastoma phenotype and have been reported to be dependent upon on Wnt signaling for proliferation [19]. IMR-32 cells were treated with BMAA over a period of 48 hours and cDNA synthesis was performed with extracted RNA. A TaqMan array containing assays for a variety of cancer-related genes was used to assess cDNA prevalence. After qPCR analysis those genes most significantly upregulated included the tumor suppressor genes PTEN and CDKN2A, Wnt pathway genes WNT1, DVL1, FZD1, and TGFBR1 (Fig 1). Functional analysis of upregulated genes was performed using the GO (gene ontology) database and PANTHER overrepresentation test, a database that provides insight into potential biological functions impacted based on the pool of genes entered. Our results linked various genes to the Wnt signaling pathway and EMT, supporting the hypothesis that BMAA may influence cancer development through its effects on Wnt signaling and cell differentiation state.

**Fig 1.**
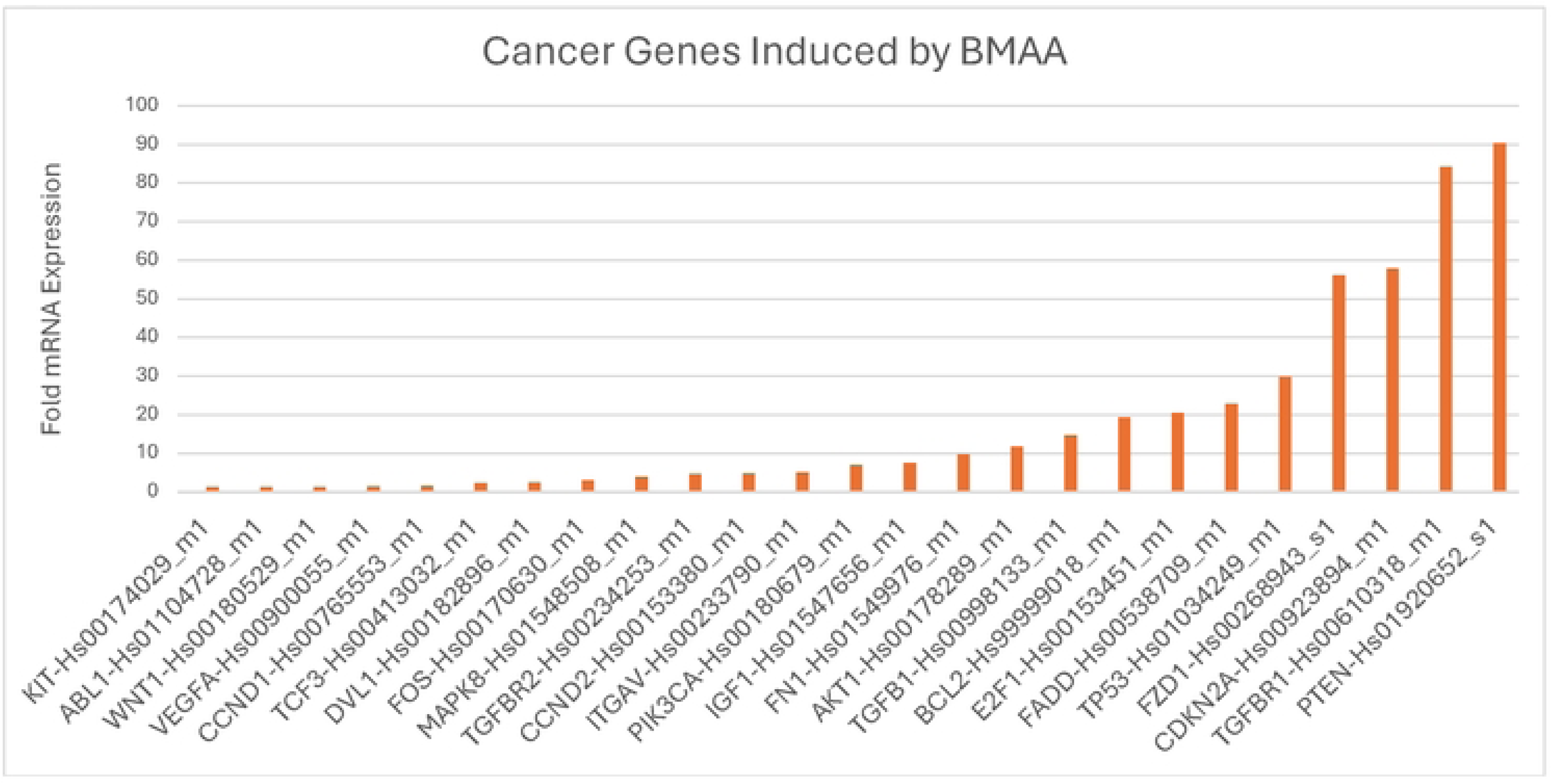
BMAA induces expression of cancer-related genes. IMR-32 cells treated with 4uM BMAA were evaluated for expression of cancer related genes via qPCR. Fold expression vs untreated cell is shown. Functional analysis of results was performed using GO ontology database and PANTHER overrepresentation test.

### BMAA enhances single cell survival

Cancer cells are characterized by resistance to apoptosis, and subsequently, survival and growth at distant areas of the body. Thus, one hallmark of mesenchymal-like cancer cells is their ability to survive as single cells and induce metastasis [20]. Wnt has been well documented as a regulator in processes that promote cancer cell development [18]. Due to BMAA’s role in mis-regulation of Wnt and promotion of cancer through EMT genes, we wondered whether BMAA could promote the EMT phenotype of increased clonal survival. IMR-32 cells were cultured in a monolayer with several sub-excitotoxic doses of BMAA over a period of 36 hours. Compared with untreated cells, cells treated with BMAA demonstrated significantly increased colony forming ability, with the largest dose exhibiting greatest clonal survival (Fig 2A, quantified in Fig 2B). These findings support the hypothesis that BMAA may influence cancer metastasis by promoting the mesenchymal-like phenotype of single cell survival.

**Fig 2.**
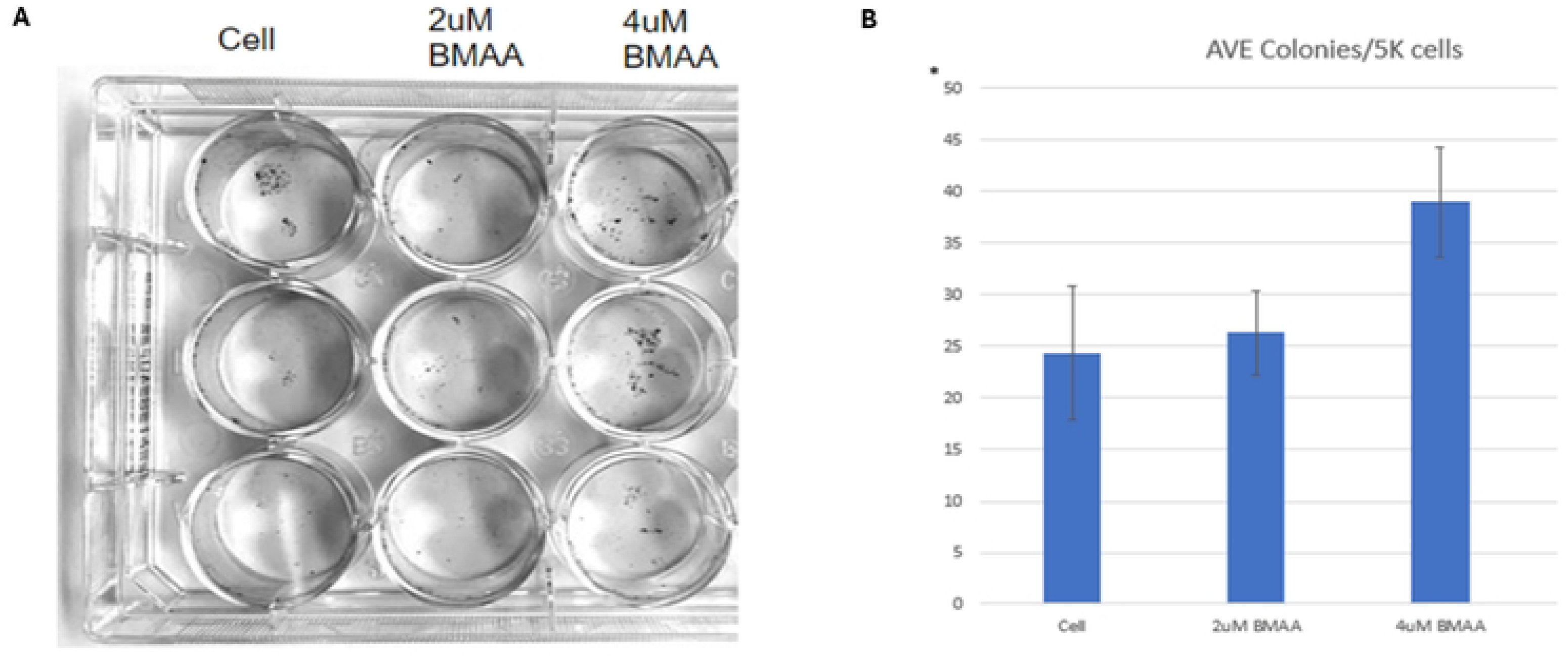
BMAA enhances single cell survival. Treatment of IMR-32 neuroblastoma cells with BMAA resulted in increas*ed colony formation. IMR-32 cells were cultured as a 2D (monolayer) culture for 36 hours in the presence of increasing concentrations of BMAA, then single cell survival was assessed by measuring the abundance of single cell colonies. Results are shown for 2uM and 4uM BMAA treatment and displayed as colony count per 5k cell.

### BMAA increases cell motility

Cancer cell metastasis relies on motility, a defining trait of mesenchymal-like cancer cells in which the cell detaches from a primary tumor and migrates to other areas of the body [20]. Dysregulation of many pathways, including Wnt signaling, play a role in acquisition of cell migration abilities [18]. We wondered then whether the presence of BMAA could influence motility in a manner similar to Wnt activation. IMR-32 cells were cultured at high density in the presence of either DMSO, BMAA, or the small molecule CHIR99021 (CHIR). CHIR is a GSK3-beta inhibitor which results in activation of Wnt/Beta-catenin signaling via stabilization of Beta-catenin. DMSO is a vehicle control for Chiron. After 48 hours, a scratch was implemented and cells were serum-starved for an additional 48-hour period (Fig 3A). In the absence of serum, little to no proliferation of cells into the scratched area is expected, therefore scratch closure is presumed to be dependent on cell motility. Treatment with BMAA and CHIR showed a significant average percent decrease in scratch width, indicating that BMAA and Wnt likely play a role in promoting the mesenchymal phenotype of cell motility (Fig 3B).

**Fig 3.**
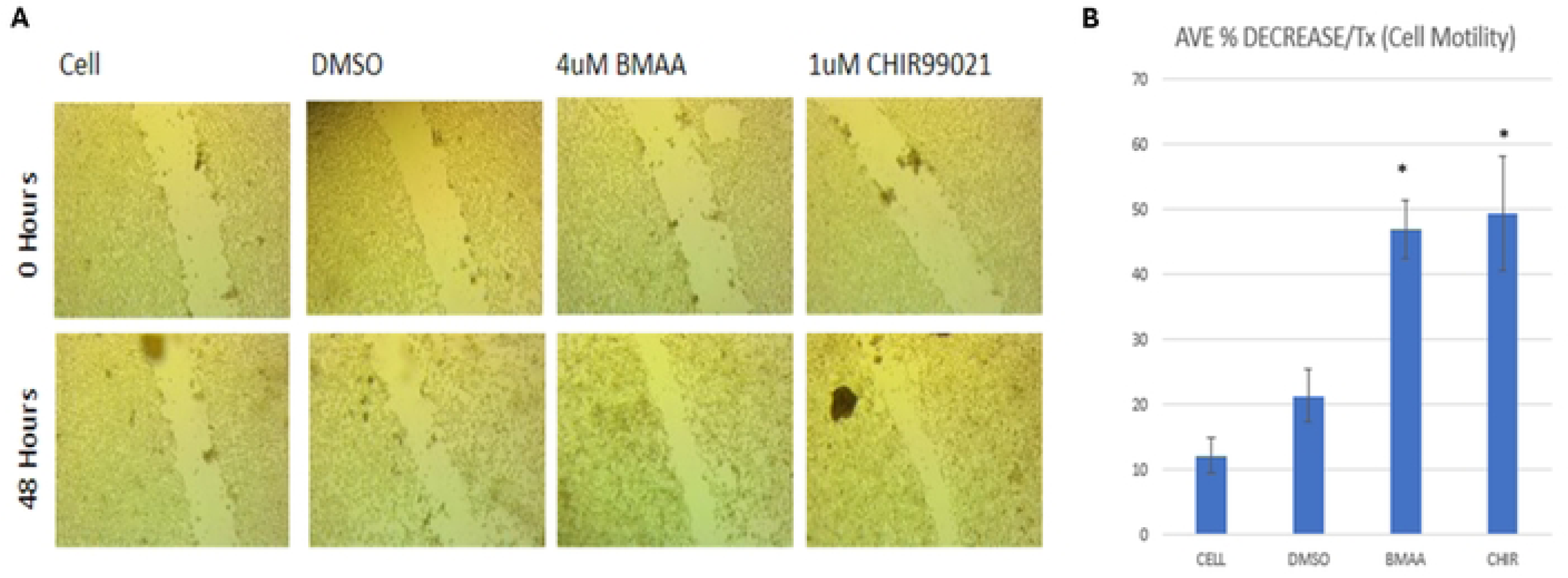
BMAA increases cell motility. Treatment of IMR-32 neuroblastoma cells with BMAA with subsequent removal of growth media resulted in increased cell motility. Similar results occurred when treated with Chiron and growth media following the same protocol. IMR-32 cells were cultured as a 2D (monolayer) culture for 48 hours in the presence of either DMSO, BMAA, or Chiron all with cell growth media. Cell media was removed and cell motility was assessed through scratch assay analysis measuring the width of the scratch. Results are shown for DMSO, 4uM BMAA, and Chiron between 0 and 48 hours.

### BMAA increases tumorshpere size

When cultured at high density over a duration of time, IMR-32 neuroblastoma cells form spheres of non-adherent or loosely adherent cells that we refer to as tumorspheres (Fig 4A). It has been documented that the cells within these tumorspheres exhibit expression of genes different from the monolayer culture and are enriched in mesenchymal-like cells [21]. Thus, we tested the effects of BMAA treatment in tumorshperes to observe the effect of the toxin on mesenchymal cell proliferation. IMR-32 cells were cultured as a monolayer for seven days to allow for tumorshpere formation. Over a period of 36 hours, tumorspheres were treated with various concentrations of BMAA. Average tumorsphere diameter and number were quantified using ImageJ analysis of micrographs for each treatment (Fig 4B and 4C). The results showed that while the presence of BMAA decreased overall average sphere number, it significantly increased tumorshpere size. This indicates that while BMAA may not be inducing the formation of new mesenchymal-like cells, it may improve the survival and/or proliferation of existing mesenchymal-like cells within these tumorspheres.

**Fig 4.**
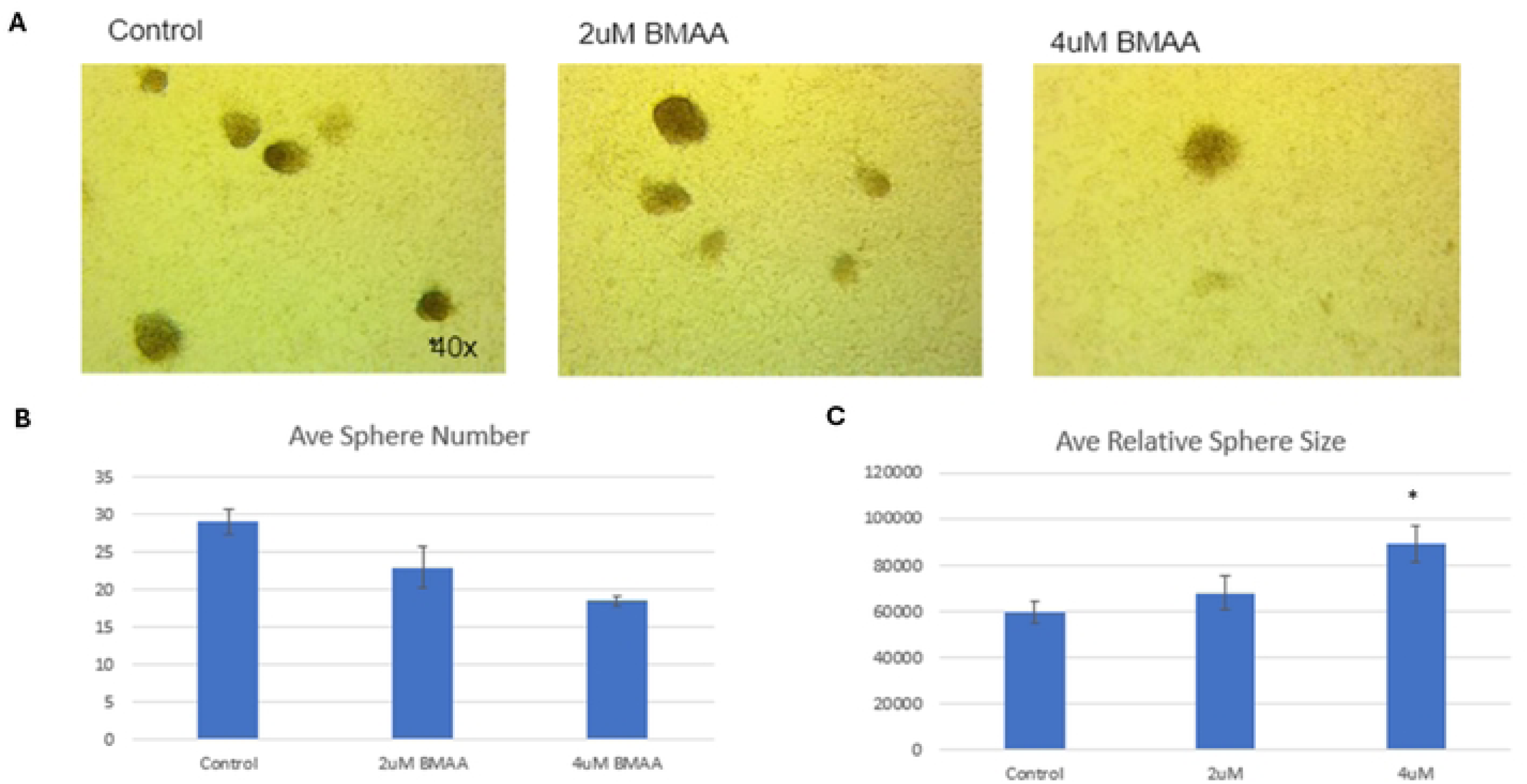
BMAA increases tumorsphere size. Treatment of IMR-32 neuroblastoma cells with BMAA resulted in increased size and decreased number of tumorspheres. IMR-32 cells were cultured as a 2D (monolayer) for 36 hours in the presence of increasing concentrations of BMAA, then proliferation was assessed by measuring the number and longest cross width diameter of the spheres using the computer program ImageJ. Results are shown for control, 2uM, and 4uM BMAA treatment.

### BMAA increases resistance to chemotherapeutics

Children that become high-risk patients of neuroblastoma typically undergo a combination of surgery and chemotherapeutics to remove the tumor [3]. Cells with a mesenchymal-like phenotype exhibit increased drug efflux pumps and anti-apoptotic effects allowing for increased resistance to cancer drugs [22]. Doxorubicin (Dox) is a commonly used anthracycline drug in neuroblastoma treatment which induces apoptosis via inhibition of topoisomerase II during DNA replication. We aimed to test the effects of BMAA on this chemotherapeutic and determine if it conferred the phenotype of cancer drug resistance associated with mesenchymal-like cancer cells [23].

Determining the effects of BMAA on Dox may also illuminate the effectiveness of this specific chemotherapeutic in improving prognosis of patients exposed to BMAA. IMR-32 cells were cultured under normal conditions with increasing concentrations of BMAA and Dox. After a period of 36 hours an LDH assay was performed to assess cell death/cytotoxicity (Fig 5). BMAA co-treatment resulted in significantly reduced cytotoxicity levels, indicating that BMAA may play a protective role against the chemotherapeutic mechanisms of Dox and promote chemoresistance. To verify the effect of BMAA on the relationship between EMT and cancer drug treatment this process was repeated for IMR-32 cells cultured as tumorspheres. Interestingly, BMAA treatment showed an even greater reduction in cytotoxicity compared to cells cultured as a monolayer. These findings suggest that the toxin may promote chemoresistance through its induction of mesenchymal-like cell proliferation.

**Fig 5.**
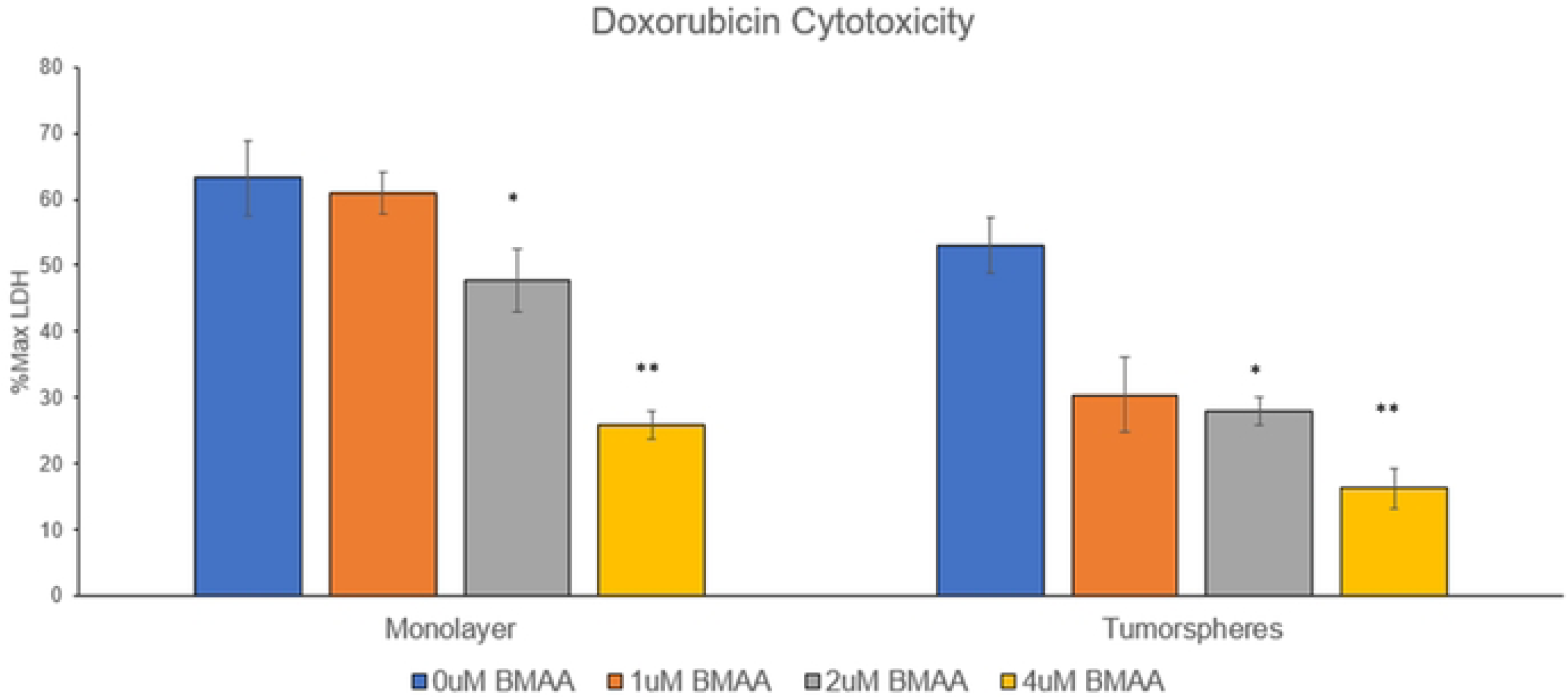
BMAA increases resistance to Doxorubicin. Co-treatment of IMR-32 neuroblastoma cells with BMAA and the chemotherapeutic Doxorubicin resulted in reduced cytotoxicity and apoptosis. IMR-32 cells were cultured as a 2D (monolayer) or 3D (tumorsphere) culture for 36 hours in the presence of increasing concentrations of BMAA and Doxorubicin, then cytotoxicity was assessed by measuring the abundance of extracellular LDH protein. Results are shown for 0.5ug/mL Doxorubicin treatment and displayed as percent of maximum LDH released from a positive control for cell lysis.

### Wnt is activated by BMAA

Prior studies have linked acute BMAA exposure to upregulation of the Wnt signaling pathway [11]. Because activation of canonical Wnt signaling is known to play a causative role in cancer cell EMT, we wondered by what mechanism BMAA may alter Wnt signaling in neuroblastoma. IMR-32 cells were cultured under normal conditions in the presence of BMAA over a period of 24 hours. RNA was collected and used to assay expression of a panel of Wnt pathway genes. The Wnt assay results showed a significant increase in expression of MYC, a downstream target of the canonical Wnt pathway that is known to play a role in regulating rapid cell division and differentiation state of cells (Fig 6). Various metalloproteinases (MMPs), a group of enzymes that function in protein degradation, were also found to be upregulated in the presence of BMAA (Fig 6). Numerous studies have tied increased MMP presence to malignant tumor development and EMT, as they are vital in degrading extracellular matrix barriers for cell migration and invasion [24]. SOX9 is another Wnt gene found to be strongly upregulated with BMAA treatment (Fig 6). Evidence suggests that increased levels of Sox9 promote stemness and survival of various cancer cell lines [25, 26]. Together, upregulation of these various Wnt genes in the presence of BMAA support the toxin’s role in mis-regulation of Wnt signaling and promotion of mesenchymal-like qualities via induction of several critical genes of the Wnt signaling pathway.

**Fig 6.**
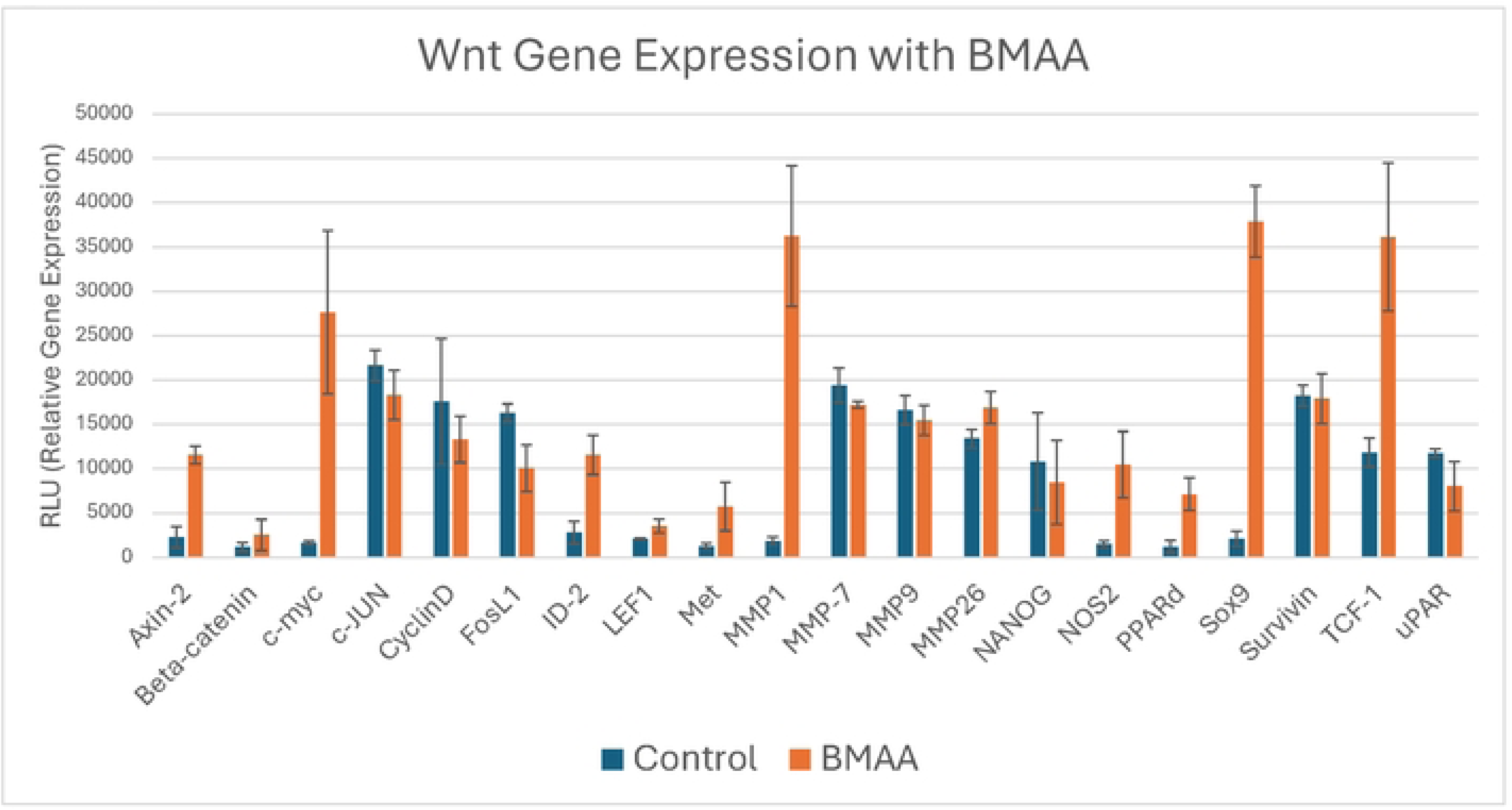
Expression of some Wnt pathway genes and Wnt targets are induced by BMAA. IMR-32 cells were treated with 4uM BMAA overnight. mRNA was extracted and used in a reverse transcription reaction to synthesize cDNA. Relative cDNA abundance was quantified via probe hybridization using chemiluminescence detection.

### BMAA-treated cells are sensitive to Myc inhibition

Myc is an oncogene involved in a large number of human cancers. Its role in contributing to essentially all hallmarks of cancer and the fact that it is a downstream target of the Wnt pathway makes it an appealing potential target for chemotherapy [27]. KJ-Pyr9 is a small molecule inhibitor of Myc/Max protein dimerization shown in prior studies to exhibit *in vivo* capability [28]. To assess the effectiveness of this inhibitor, IMR-32 cells were cultured under normal conditions in the presence of KJ Pyr9 over a period of 48 hours. Higher concentrations of KJ Pyr9 correlated with a decrease in proliferation, as measured by XTT assay, indicating that it is an effective inhibitor of rapid cell growth (Fig 7A). Of note, extracellular LDH, an indicator of cell lysis, was not increased by KJ-Pyr9, indicating that the Myc inhibitor did not induce apoptosis. These results point to KJ Pyr9 effectively arresting cell division without inducing cell death. We then wondered if the Myc inhibitor would exhibit similar activity in the presence of BMAA. IMR-32 cells were cultured under normal conditions in the presence of increasing concentrations of KJ Pyr9 and BMAA over a period of 48 hours. Proliferation was found to significantly decrease in IMR-32 cells exposed to KJ Pyr9 in doses of 1.25uM or higher when compared to normal conditions (Fig 7B). Interestingly, cells co-treated with the Myc inhibitor and BMAA displayed no significant difference to controls, with the impact of BMAA on proliferation decreasing with higher doses of KJ Pyr9 (Fig 7B). Together, these results suggest that KJ Pyr9, unlike Doxorubicin (Fig 5), is capable of arresting cell division even in the presence of BMAA.

**Fig 7.**
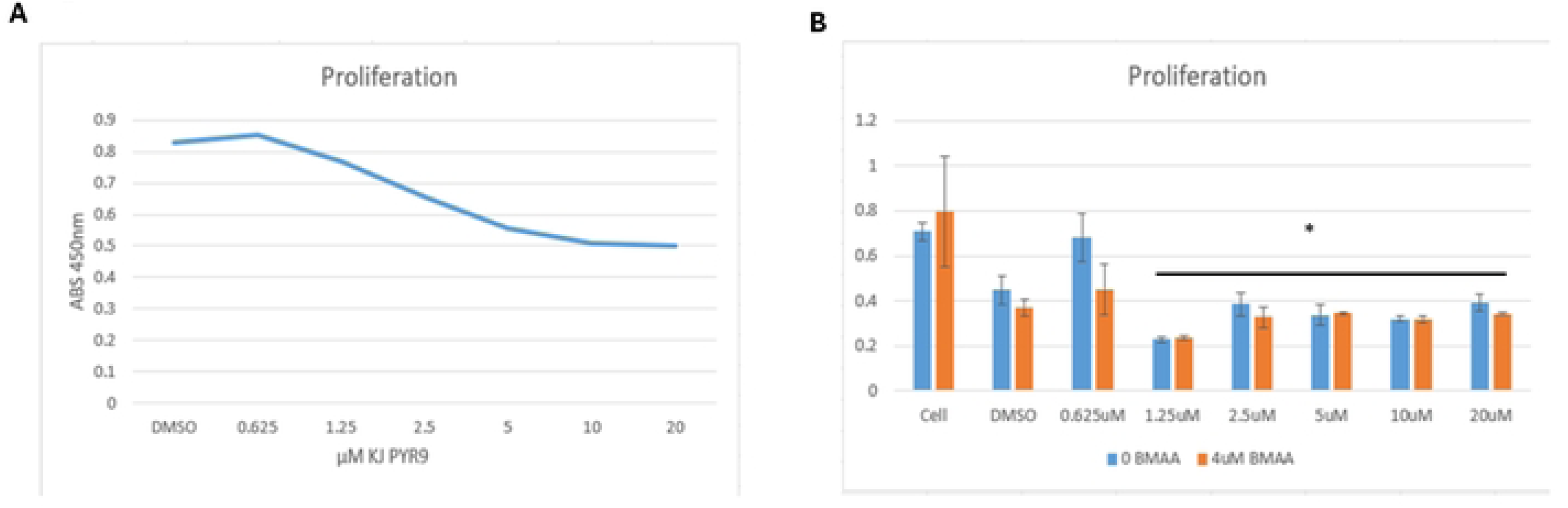
BMAA treated cells are sensitive to Myc inhibition. Co-treatment of IMR-32 neuroblastoma cells with BMAA and various concentrations of a Myc inhibitor (KJ Pyr9) showed a decrease in proliferation without affecting cytotoxicity. IMR-32 cells were cultured as a 2D (monolayer) culture for up to 48 hours in the presence of varying concentrations of BMAA and Myc inhibitor, then proliferation was assessed using a colorimetric tetrazolium assay. Results are shown for 0.625uM to 20uM Myc inhibitor treatment and displayed as absorbance 450. No significant difference was observed between cells treated or not treated with BMAA.

### BMAA’s effect on tumorspheres is Myc-dependent

Our prior experiments validated the role of BMAA in promoting proliferation of mesenchymal-like cells in tumorspheres (Fig 4). We then wondered if this relationship was Myc dependent, as BMAA is an apparent up regulator of Wnt signaling. IMR-32 tumorspheres treated with BMAA alone over a period of seven days exhibited a significant increase in fold area change (Fig 8A, quantified in Fig 8B). Conversely, spheres co-treated with BMAA and XAV939, a Wnt inhibitor, showed no increase in fold area change at day seven when compared to normal conditions (Fig 8A, quantified in Fig 8B). Together these results support the hypothesis that BMAA influences proliferation of tumorsphere cells in a Wnt-dependent manner. To elucidate whether this phenomenon is also Myc-dependent, IMR-32 cells were co-treated with BMAA and KJ-PYR9, a Myc inhibitor. These tumorshperes experienced a significant decrease in fold area change, indicating that the effect of BMAA on tumorshpere proliferation is in fact dependent upon Myc activity (Fig 8A, quantified in Fig 8B). Importantly, the finding that BMAA-induced tumorsphere growth could be reversed by Myc inhibition further suggests the therapeutic potential of a Myc-inhibitor in neuroblastoma treatment. Although different growth rates of spheres were observed between conditions, all treatments except the myc inhibitor exhibited irregularities in cell morphology over the seven-day period (Fig 8A). We hypothesize that this effect may be a result of decreased control over cell patterning and physical instability of larger spheres. Further research is needed to properly assess the mechanism behind these morphological irregularities.

**Fig 8.**
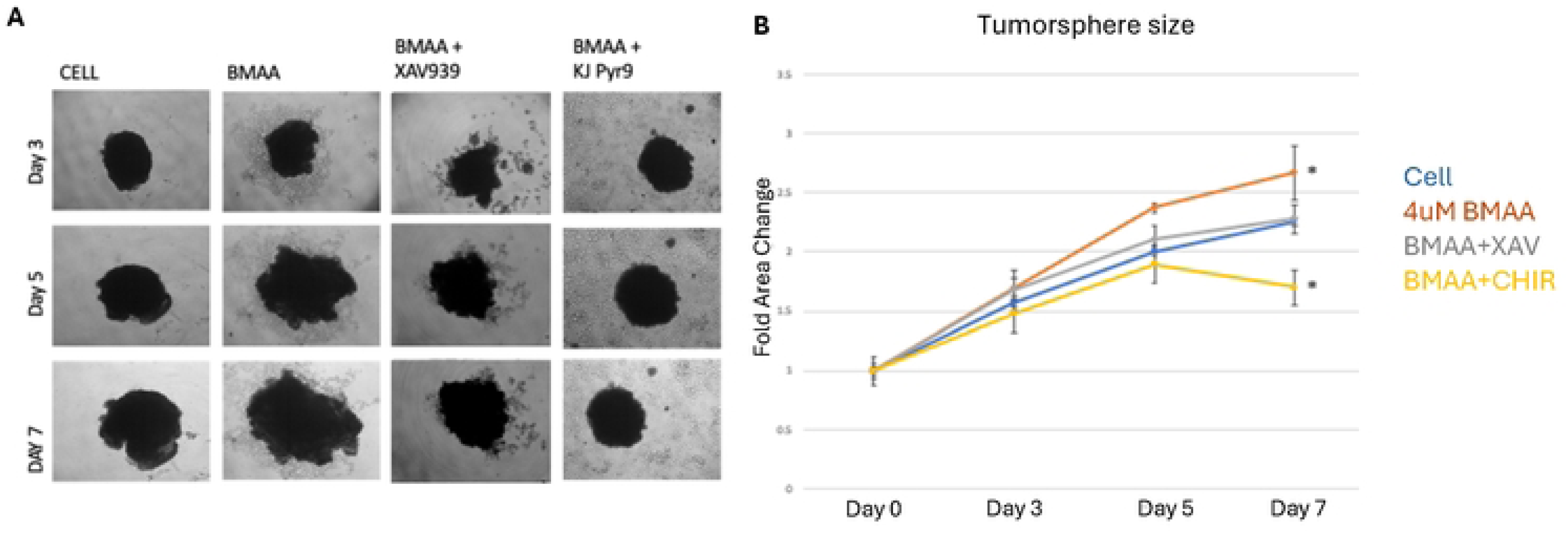
BMAA does not protect against Myc inhibition. Treatment of IMR-32 neuroblastoma cells with BMAA increased cell proliferation compared to untreated cells. Co-treatment with BMAA and a Wnt Inhibitor (XAV939) resulted in similar growth as untreated cells, whereas co-treatment with BMAA and a Myc inhibitor (KJ Pyr9) resulted in decreased cell growth. IMR-32 cells were cultured as a 3D (tumorsphere) for 7 days in the presence of increasing concentrations of BMAA, then sphere area was quantified using ImageJ analysis software. Results are shown for BMAA, Myc inhibitor, and Wnt inhibitor, measured in triplicates and taking the average area at time points 0, 3, and 7 days.

### BMAA enhances cell invasion

Chemotaxis, in the context of cancer, is the organ-specific migration of cells in response to an extracellular gradient. Cancer cell metastasis and survival relies on dissemination, invasion, and growth at distant sites throughout the body. Various factors such as metalloproteinase expression, chemotaxis, single cell survival, and cell motility play a large role in these crucial steps [29]. To assess these aspects of metastasis, IMR-32 cells were plated in a matrigel transwell assay, in which cells are serum starved and placed above cell media with a porous membrane in between permitting chemotaxis. For cells to survive, they must invade the Matrigel and digest their way into the lower chamber containing serum. Thus, invasion is an indicator of metastatic potential, another characteristic of mesenchymal-like cancer cells. IMR-32 cells treated with the Wnt inhibitor, XAV939, showed a significant decrease in invasion; whereas cells treated with a Wnt activator, CHIR99021, exhibited a significant increase in invasive potential (Fig 9). These results support the hypothesis that Wnt activity plays a key role in regulating cancer cell invasion. Cells treated with a Myc inhibitor showed a large decrease in invasion, indicating that Wnt regulates cancer cell invasion in a Myc-dependent manner (Fig 9). Treatment of IMR-32 cells with BMAA alone showed an increase in invasion, highlighting the potential of BMAA to promote metastatic potential via induction of Wnt (Fig 9). Interestingly, cells co-treated with BMAA and either XAV939 or KJ Pyr9 invasion were found to decrease invasiveness, demonstrating that the effects of BMAA on cancer cell invasion can be arrested through inhibition of Wnt and Myc activity (Fig 9).

**Fig 9.**
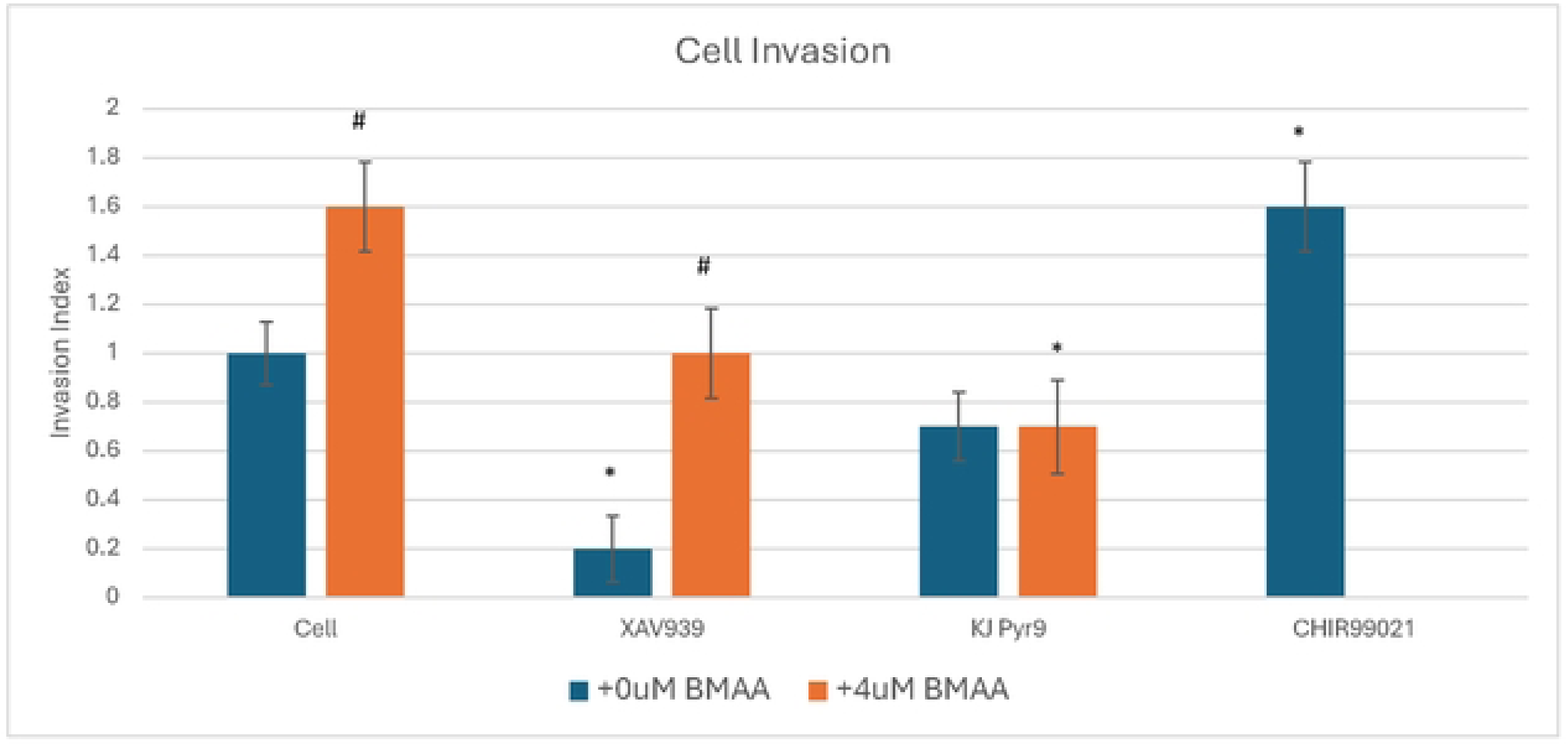
BMAA enhances cell invasion. IMR-32 cells were plated in serum-free media in a transwell plate and allowed to migrate toward growth media in the lower chamber. Migrating cells were quantified by staining the lower side of the transwell membrane after 48 hours. Data shown is presented as the invasion index (compared to untreated cell control). CHIR99021 was used as a positive control for Wnt activation (no BMAA+CHIR99021 treatment was performed). * = vs. cell control, # = vs. 0uM BMAA

### Conclusions

Together our findings suggest that BMAA may promote aggressive neuroblastoma through its effects on cell epithelial to mesenchymal transition, specifically through Wnt/Myc activation. BMAA appears to promote various EMT hallmarks such as increased single cell survival, motility, proliferation, and resistance to chemotherapeutics, as well as upregulate genes associated with EMT including Wnt targets and others such as TGFBR1 [30, 31]. Due to the involvement of EMT in tumor progression and metastasis, our results suggest that while BMAA has been extensively tied to neurodegeneration, it may also play a role in cancer development, progression, and relapse.

BMAA is a known mis-regulator of canonical Wnt signaling, however, the mechanism through which it exerts these effects is not fully studied [11]. The present study reveals an increase in expression of the Wnt target gene MYC and other Wnt pathway genes related to differentiation and metastasis in the presence of BMAA, further supporting its role in cell de-differentiation through Wnt dysregulation. Some drugs directly targeting Wnt have shown various off-target effects, indicating that targeting downstream molecules of Wnt may provide a better alternative [32]. MYC is an oncogene involved in many cancers. Its widespread cancer relevance and placement as a downstream target of Wnt makes this gene an appealing potential target for chemotherapy [27, 28]. Our results showed that while a commonly used chemotherapeutic in neuroblastoma treatment, Doxorubicin, was ineffective in the presence of BMAA, cells remained sensitive to growth inhibition induced by a Myc inhibitor [23]. With this information, treatment plans could be adapted to combat the effects of BMAA and mesenchymal transition in neuroblastoma. Using Myc inhibition in conjunction with, or in place of Doxorubicin, treatments could potentially improve the prognoses of children with high-risk Neuroblastoma and decrease the opportunity for relapse driven by mesenchymal cancer cell proliferation.

Being an environmental toxin, an escalation of climate change could pose serious health issues related to BMAA. As temperatures rise, warmer water creates ideal conditions to expand the range of algae blooms and increase human exposure to BMAA [15]. Additionally, changes in precipitation patterns and increased frequency of extreme weather events associated with climate change can lead to nutrient runoff from agricultural areas, urban landscapes, and industrial sites into water bodies. These nutrients, particularly nitrogen and phosphorus, fuel the growth of the primary BMAA producer, cyanobacteria [33]. Together, these events could cause more people to face the effects of BMAA, highlighting the importance of understanding what role BMAA plays in human health.

## Funding and additional information

Research reported in this publication was supported by the South Dakota Biomedical Research Infrastructure Network (SD BRIN) through an Institutional Development Award (IDeA) from the National Institute of General Medical Sciences of the National Institutes of Health under grant number P20GM103443. The content is solely the responsibility of the authors and does not necessarily represent the official views of the National Institutes of Health. Cell culturing and functional assays were performed by WestCore at Black Hills State University. The funders had no role in study design, data collection and analysis, decision to publish, or preparation of the manuscript.

## Conflict of Interest

"The authors declare that they have no conflicts of interest with the contents of this article."

## Notes

### Competing Interest Statement

The authors have declared no competing interest.

